# The multistep quality control of mitochondrial β-barrel protein import

**DOI:** 10.64898/2026.05.24.727513

**Authors:** Elena T. Kämmerer, Nicole Okon, Niklas Limbach, Janet Hinnenkamp, Enzo Scifo, Gesa Schmidt, Frederike Eiding, Lara Calvo Santos, Max-Hinderk Schuler, Thomas Becker, Fabian den Brave

**Affiliations:** Institute of Biochemistry and Molecular Biology, University Hospital Bonn, University of Bonn, 53115 Bonn, Germany; Institute of Biochemistry and Molecular Biology, Medical Faculty, University of Freiburg, 79104 Freiburg, Germany

**Author notes:** Gene center and Department of Biochemistry, Ludwig-Maximilians-University of Munich, 81377 Munich, Germany (Max-Hinderk Schuler).

**Keywords:** MitoStores, mitochondria, outer membrane, protein degradation, mitoTAD, protein translocase, SAM complex, TOM complex

## Abstract

Mitochondrial β-barrel proteins fulfill essential functions for cell viability including transport of proteins, ions and metabolites across the outer membrane. The translocase of the outer membrane (TOM complex) and the sorting and assembly machinery (SAM complex) transport these proteins into the outer membrane. Although β-barrel proteins are essential for cell viability, the quality control mechanisms that monitor their import into the outer membrane remain unknown. We uncovered that Msp1 and Ubx2-recruited Cdc48 cooperate to remove a SAM-arrested Tom40 variant. Prior to degradation, the β-barrel protein is deposited in cytosolic protein storages, termed MitoStores, to minimize accumulation of aberrant proteins in the cell. The cytosolic AAA ATPase Hsp104 disaggregates these proteins to facilitate their proteasomal degradation. Altogether, mitochondrial and cytosolic factors cooperate to degrade translocation-arrested mitochondrial β-barrel proteins.

## INTRODUCTION

Mitochondria emerged by an endosymbiotic event 2 billion years ago. One relic of the endosymbiotic origin are proteins with a transmembrane β-barrel, which exist in the outer membrane of the endosymbiosis-derived organelles, mitochondria and plastids, as well as in the outer membrane of Gram-negative bacteria ^1–3^. Mitochondria contain five β-barrel proteins in bakeŕs yeast *Saccharomyces cerevisiae* that carry out essential functions for cell viability ^2,4–6^. Tom40 forms the protein-conducting channel of the translocase of the outer mitochondrial membrane (TOM complex), whereas Sam50 (also termed Tob55) is the core subunit of the sorting and assembly machinery (SAM complex) that inserts β-barrel proteins. Por1 (also termed VDAC for voltage-depended anion channel) and its isoform Por2 form channels for the passage of ions and metabolites across the outer membrane. Finally, the β-barrel protein Mdm10 is present in two protein complexes. First, it constitutes the outer membrane anchor of the endoplasmic reticulum encounter structure (ERMES) that forms a bridge between both organelles for lipid trafficking and to maintain mitochondrial morphology ^7,8^. Second, Mdm10 binds to the SAM complex to promote assembly of TOM subunits ^8–12^.

Precursors of β-barrel proteins are produced on cytosolic ribosomes and have to be imported into mitochondria. Cytosolic chaperones guide the precursor proteins to the Tom20 and Tom70 receptors of the TOM complex ^13–15^. The Tom40 pore allows transport of these proteins across the outer membrane ^13,16–20^. Small TIM chaperones of the intermembrane space bind to the imported precursor proteins and transfer them to the SAM complex ^21–23^. TOM and SAM complexes form a supercomplex to facilitate the transfer of β-barrel proteins ^24,25^. Precursors of β-barrel proteins contain a conserved β-signal within the last β-strand, which is recognized by the SAM components ^26^. The SAM complex consists of the β-barrel protein Sam50 and the two peripherally bound Sam35 (Tob38) and Sam37 (Mas37, Tom37) ^17,18,27–32^. Sam50 forms a lateral gate between the first and the last β-strand in which the incoming precursor is inserted in a stepwise manner ^32–35^. Once the β-barrel is folded, it is released into the outer membrane. The SAM complex also promotes the assembly of Tom40 with small Toms to initiate the first steps in the formation of the TOM complex ^36–39^. Mdm10 cooperates with the SAM complex to release the Tom40-small Toms intermediate into the outer membrane and to promote its assembly with Tom22 to form the TOM complex ^8–11^.

A network of quality control components monitors protein import into mitochondria and removes non-imported mitochondrial proteins ^40–44^. Import defects can occur upon premature folding of the precursor proteins since only largely unfolded proteins can pass the TOM channel. Furthermore, damage of the protein translocase or depletion of the mitochondrial membrane potential can impair protein import into mitochondria ^41,45–50^. The accumulation of non-imported precursor proteins in the cell causes proteotoxic stress and eventually cell death ^45,51,52^. The cells have implemented several strategies to minimize proteotoxic stress. Non-imported precursor proteins are distributed to different cellular compartments and are degraded by the proteasome ^45,51–54^. Several of the non-imported mitochondrial precursor proteins can be sequestered in protein deposits, termed MitoStores ^55–57^. Recent studies have uncovered quality control mechanisms that remove precursor proteins that arrest at the TOM complex during import ^46,58,59^. Two AAA-ATPases exert central functions in these processes. The outer membrane-bound Msp1 extracts precursors from the TOM complex under stress conditions in the mitochondrial compromised protein import response (mitoCPR) ^47,58^, whereas Ubx2 recruits the cytosolic AAA-ATPase Cdc48 that removes precursor proteins under constitutive conditions in the mitochondrial protein translocation-associated degradation (mitoTAD) ^46^. So far, the identification of quality control factors has been limited to presequence-containing precursor proteins. How transport-arrested mitochondrial proteins lacking a cleavable presequence are degraded remains largely unknown.

While the integration of β-barrel proteins into the outer membrane is well understood, the quality control mechanisms that remove unproductive import intermediates remain unclear. Here, we studied the degradation of a Tom40 variant that accumulates at the SAM complex. This Tom40 variant is ubiquitylated and rapidly degraded by the proteasome. Using deletion mutants, we could show that Msp1 extracts the Tom40 variant to facilitate its ubiquitylation. Ubx2 recruits the cytosolic AAA ATPase Cdc48 to complete removal of the ubiquitylated β-barrel protein from the outer membrane. Outside mitochondria, the aberrant Tom40 variant is sequestered in protein deposits, the MitoStores, to minimize proteotoxicity for the cell. The AAA ATPase Hsp104 disaggregates this protein to promote its proteasomal degradation. Altogether, the degradation of SAM-arrested β-barrel proteins is a multi-step process involving three AAA ATPases Msp1, Cdc48 and Hsp104, transient deposition in the MitoStores and proteasomal degradation.

## RESULTS

### Tom40-18 is an in vivo clogger of the SAM complex

In order to study degradation of translocation-arrested β-barrel proteins, we screened our library of temperature-sensitive Tom40 mutant strains for a Tom40 variant that accumulates at the SAM complex. Cells expressing the *TOM40-18* variant display a temperature-sensitive growth defect (Figure 1A, B) ^60^. We isolated mitochondria from cells grown under permissive conditions and found that the steady state levels of Tom40 were mildly decreased in the mutant mitochondria, while other TOM subunits and outer membrane proteins remain largely unchanged (Supplemental Figure 1A). To assess the integrity of outer membrane translocases, mitochondria were solubilized with the non-ionic detergent digitonin and analyzed by blue native electrophoresis. The TOM complex co-migrating with the 440 kDa size marker was strongly reduced in *tom40-18* mitochondria when detected with antisera against Tom22 and Tom40 (Figure 1C, lanes 1-4), indicating that the formation of the TOM complex is impaired in the mutant mitochondria. The SAM complex exists in wild-type mitochondria in two major populations that can be resolved by blue native electrophoresis: The SAM_core_ complex consisting of Sam50, Sam35 and Sam37 and the SAM-Mdm10 complex that additionally contains Mdm10 (Figure 1C, lane 6) ^8,9,34^. In *tom40-18* mitochondria, the levels of both SAM_core_ and SAM-Mdm10 complexes were markedly reduced (Figure 1C, lanes 5-8). However, a larger SAM* is strongly increased that does not correspond to SAM-Mdm10 (Figure 1C, lane 5). Instead, the SAM* form represents the SAM complex with bound Tom40 (Becker et al., 2011; Taneka et al., 2021). Supporting this conclusion, immunodetection with a Tom40-specific antiserum revealed a band co-migrating with SAM* (Figure 1C, lane 4). We next asked whether accumulation of Tom40-18 at the SAM complex interferes with the import of β-barrel proteins. Therefore, we synthesized ^35^S-labelled Tom40 and Por1 in cell-free translation extracts and imported both β-barrel proteins into isolated *tom40-18* mitochondria and analyzed the import reaction by blue native electrophoresis. Por1 assembles into high molecular weight protein complexes ^13,61^, which was largely impaired in *tom40-18* mutant mitochondria (Figure 1D). Tom40 import proceeds through distinct assembly intermediates that that can be resolved by blue native electrophoresis: Tom40 first binds to the SAM complex (SAM), followed by the formation of assembly intermediate II that contains the small Tom proteins and Tom40 (Int. II) and the final integration into the mature TOM complex (TOM). In *tom40-18* mitochondria, association of newly imported Tom40 with the SAM complex was already impaired (Figure 1D). As control, we imported radio-labeled precursors of Atp2 and Su9-DHFR in *tom40-18* mitochondria (Figure 1E). Non-imported precursor proteins were removed by proteinase K. Although the TOM complex levels were reduced, the import of Atp2 and Su9-DHFR in *tom40-18* mitochondria was only mildly affected (Figure 1E), excluding a general import defect in these mutant mitochondria. We conclude that the Tom40-18 variant accumulates at the SAM complex and clogs the SAM complex for further import of β-barrel proteins.

**Figure 1.**
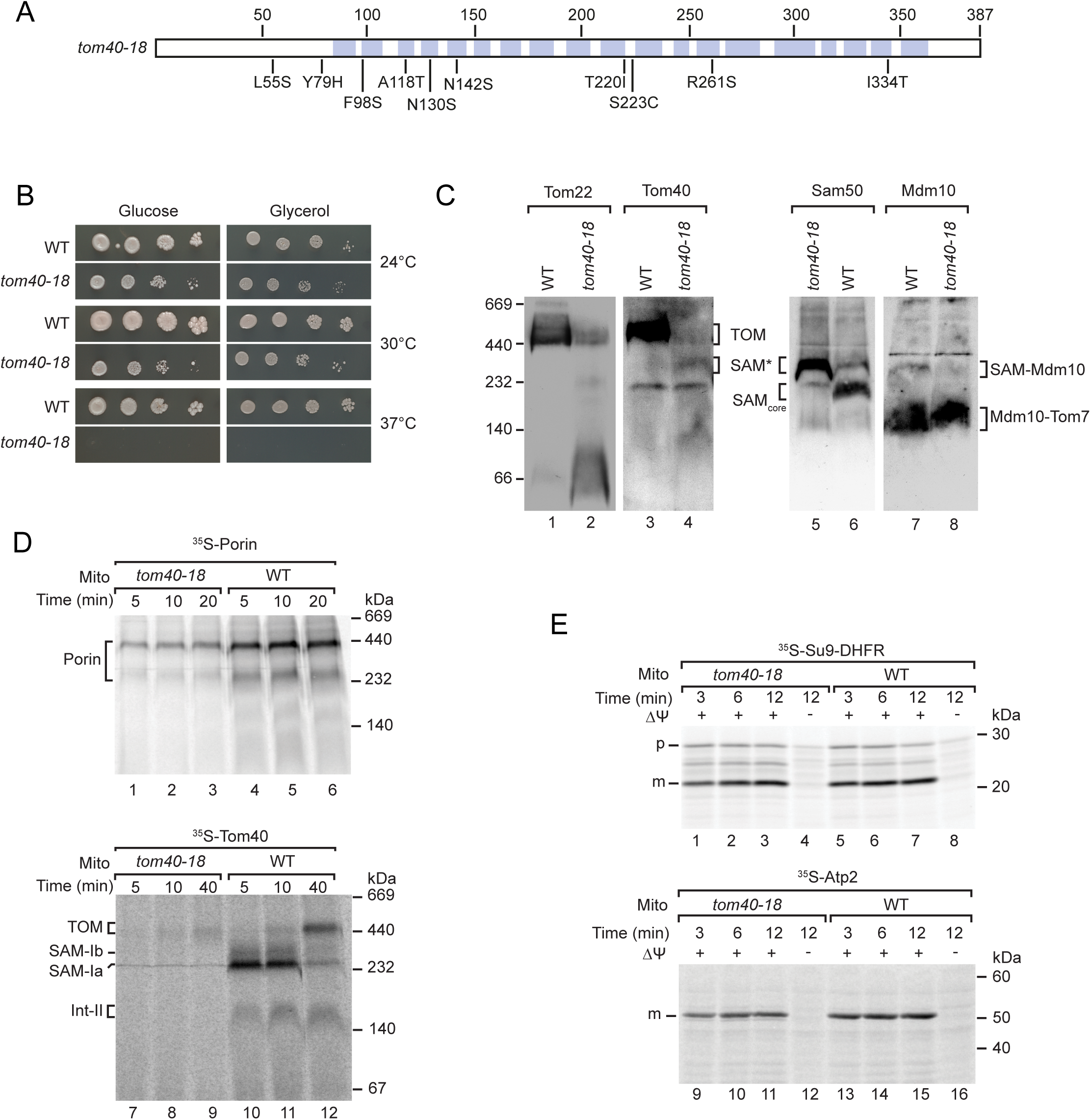
The Tom40-18 variant arrests at the SAM complex. **(A)** Schematic representation of the amino-acid changes in Tom40-18 compared to Tom40. **(B)** Serial dilutions of wild-type (WT) and *tom40-18* strains were grown on medium containing either glucose or glycerol as carbon source at the indicated temperature. **(C)** WT and *tom40-18* mitochondria were analyzed by blue native electrophoresis and immunodetection with the indicated antisera. **(D)-(E)** ^35^S-labelled Por1, Tom40 (D) or Atp2 and Su9-DHFR (E) were imported for the indicated time periods into WT and *tom40-18* mitochondria and analyzed by blue native electrophoresis (D) or SDS-PAGE (E). After import of the precursor of Atp2 and Su9-DHFR, mitochondria were treated with proteinase k to remove non-imported proteins. Where indicated, the membrane potential (Δψ) was depleted to block import.

### The Tom40-18 variant is degraded by the ubiquitin-proteasome system

We used the Tom40-18 variant to analyze how SAM-arrested β-barrel proteins are removed. To minimize pleiotropic effects, we co-expressed plasmid-borne TOM40-18 fused C-terminally to a hemagglutinin (HA)-tag in wild-type cells that contain a wild-type copy of Tom40 (Figure 2A). We used a strong ADH promoter to ensure that the Tom40-18_HA_ levels were comparable to those of endogenous Tom40. Expression of TOM40-18_HA_ impaired growth of wild-type cells, indicating that the variant is toxic for the cell (Supplemental Figure 1B). The Tom40-18_HA_ variant was targeted to mitochondria as shown by immune fluorescence using antibodies against the HA-tag (Supplemental Figure 1C). We next used carbonate extraction to assess whether Tom40-18_HA_ was integrated into the outer membrane ^37,62^. At alkaline pH, peripheral proteins are extracted from the membrane into the soluble fraction as exemplified by Tim10, whereas integral membrane proteins remain in the pellet fraction (Supplemental Figure 1D)^37^. Similar to a SAM-arrested Tom40 precursor ^24^, Tom40-18_HA_ was detected in the pellet fraction (Supplemental Figure 1D). To test whether Tom40-18_HA_ had passed the outer membrane, we treated isolated mitochondria with proteinase K. Like wild-type Tom40_HA_, the Tom40-18_HA_ was protected in intact mitochondria and the HA-tag became accessible to the protease only after osmotic rupture of the outer membrane (Supplemental Figure 1E). Finally, we tested whether Tom40-18_HA_ accumulated at the SAM complex by affinity purification utilizing the HA-tag. While the co-purification of TOM subunits was reduced compared to Tom40_HA_, we detected increased levels of Sam50 in the elution fraction (Figure 2B). We conclude that Tom40-18_HA_ accumulates at the SAM complex.

**Figure 2.**
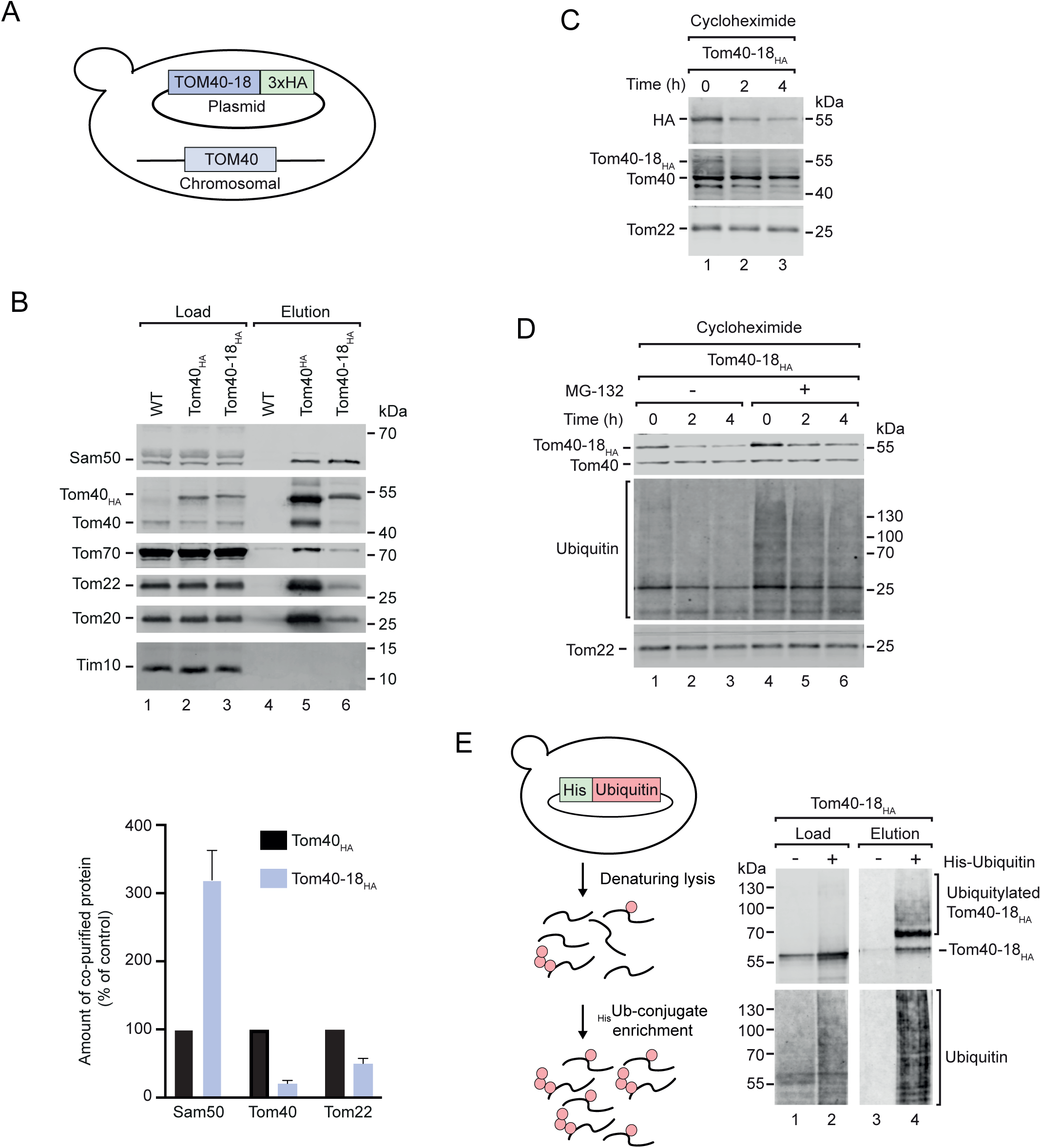
SAM-arrested TOM40-18_HA_ is degraded via the ubiquitin-proteasome pathway. **(A)** Scheme of the used strain expressing Tom40-18_HA_ variant from a plasmid. **(B)** Upper panel: Total cell extracts from wild-type (WT), Tom40_HA_ and Tom40-18_HA_ were subjected to affinity purification. Load (1%) and elution (100%) fractions were analyzed by SDS-PAGE and immunodetection with the indicated antisera. Lower Panel: Quantification of Sam50, Tom40 and Tom22 co-purified with Tom40-18_HA_ relative to Tom40_HA_. Mean of three independent experiments +/- S.E.M. **(C)** WT cells expressing Tom40-18_HA_ were treated with cycloheximide (CHX) for the indicated time periods. Cell extracts were analyzed by SDS-PAGE and immunodetection with the indicated antisera. **(D)** *pdr5Δ* cells expressing Tom40-18_HA_ were treated with cycloheximide (CHX) in the absence or presence of the proteasomal inhibitor MG132 for the indicated time periods. Cell extracts were analyzed by SDS-PAGE and immunodetection with the indicated antisera. **(E)** Left panel: Experimental setup for purification of ubiquitin-modified proteins under denaturing conditions from cells expressing His-tagged ubiquitin. Right panel: WT cells expressing Tom40-18_HA_ with or without expression of His-tagged ubiquitin were subjected to affinity purification under denaturing conditions. Load (1%) and elution (100%) fractions were analyzed by SDS-PAGE and immunodetection with the indicated antisera.

To investigate how the SAM-bound Tom40-18_HA_ is removed, we first monitored its degradation over time after blocking of protein synthesis with cycloheximide. Using a Tom40-specific antiserum, we detect both Tom40-18_HA_ and the endogenous Tom40. Tom40-18_HA_ was degraded, whereas endogenous Tom40 remained stable (Figure 2C). Treatment of the cell with the proteasome inhibitor MG132 inhibited degradation of Tom40-18_HA_ (Figure 2D), revealing that Tom40-18_HA_ was degraded by the proteasome. Proteasomal degradation typically involves ubiquitylation of the client protein ^63^. To detect ubiquitylated forms of Tom40-18_HA_, we co-expressed His-tagged ubiquitin and performed affinity purification under denaturing conditions ^59^. High-molecular weight forms of Tom40-18_HA_ were recovered in the pulldown via His-tagged ubiquitin, indicating that Tom40-18_HA_ is ubiquitylated (Figure 2E). We conclude that Tom40-18_HA_ is degraded by the cytosolic ubiquitin-proteasome system.

### Two AAA ATPases extract the Tom40-18_HA_ variant from mitochondria

We asked how Tom40-18_HA_ is removed from the outer membrane. Two AAA ATPases have been previously reported to extract proteins from the outer mitochondrial membrane. The outer membrane embedded AAA ATPase Msp1 dislocates mislocalized proteins from the outer membrane ^64–66^ and removes precursor proteins from the TOM complex ^47,58^. Ubx2 forms a docking site for the cytosolic AAA ATPase Cdc48 that extracts translocation-stalled precursor proteins from the TOM complex ^46^ and certain outer membrane proteins ^67–70^. We first investigated whether the AAA ATPases associate with Tom40-18_HA_ by affinity purification utilizing the HA-tag. Indeed, we detected Msp1, Ubx2 and Cdc48 in the elution fractions (Figure 3A, lane 4). For comparison, Sam50 and small amounts of TOM subunits were detected, whereas Mim1 was not found in the elution fraction (Figure 3A, lane 4). Having established that Tom40-18_HA_ associates with the two AAA ATPases, we asked whether they promote the removal of Tom40-18_HA_. We expressed Tom40-18_HA_ in *msp1*Δ and *ubx2*Δ strains and found that the growth of the cells was impaired compared with wild-type cells (Figure 3B), indicating that the two quality control components contribute to clearance of the SAM-arrested protein. To directly study the degradation of Tom40-18_HA_, we treated the cells with cycloheximide for different time points to inhibit ribosomal protein biosynthesis and monitor protein levels over time ^46,59^. Degradation of Tom40-18_HA_ was delayed in both *msp1*Δ and *ubx2*Δ cells compared with wild-type cells, whereas endogenous wild-type Tom40 remained stable under these conditions (Figure 3C). We therefore conclude that both Msp1 and Ubx2/Cdc48 promote extraction of Tom40-18_HA_ to facilitate its degradation. To study a possible interplay of both components, we used a *ubx2Δ msp1Δ* double deletion strain ^46^. Interestingly, the degradation pattern was largely unchanged in comparison to the single deletion strains (Figure 3C), indicating that Msp1 and Ubx2/Cdc48 cooperate in the removal of Tom40-18_HA_. Supporting this view, we found less ubiquitylation of Tom40-18_HA_ in the absence of Msp1, while upon deletion of *UBX2* ubiquitylated Tom40-18_HA_ accumulated (Figure 3D). We conclude that Msp1 extracts Tom40-18_HA_ to facilitate ubiquitylation. Ubx2 contains a UBA domain that binds ubiquitylated proteins and a UBX domain that forms a docking site for Cdc48 ^46,71,72^. Thus, Ubx2 recruits Cdc48 to ubiquitylated Tom40-18_HA_ to complete the extraction of the β-barrel protein.

**Figure 3.**
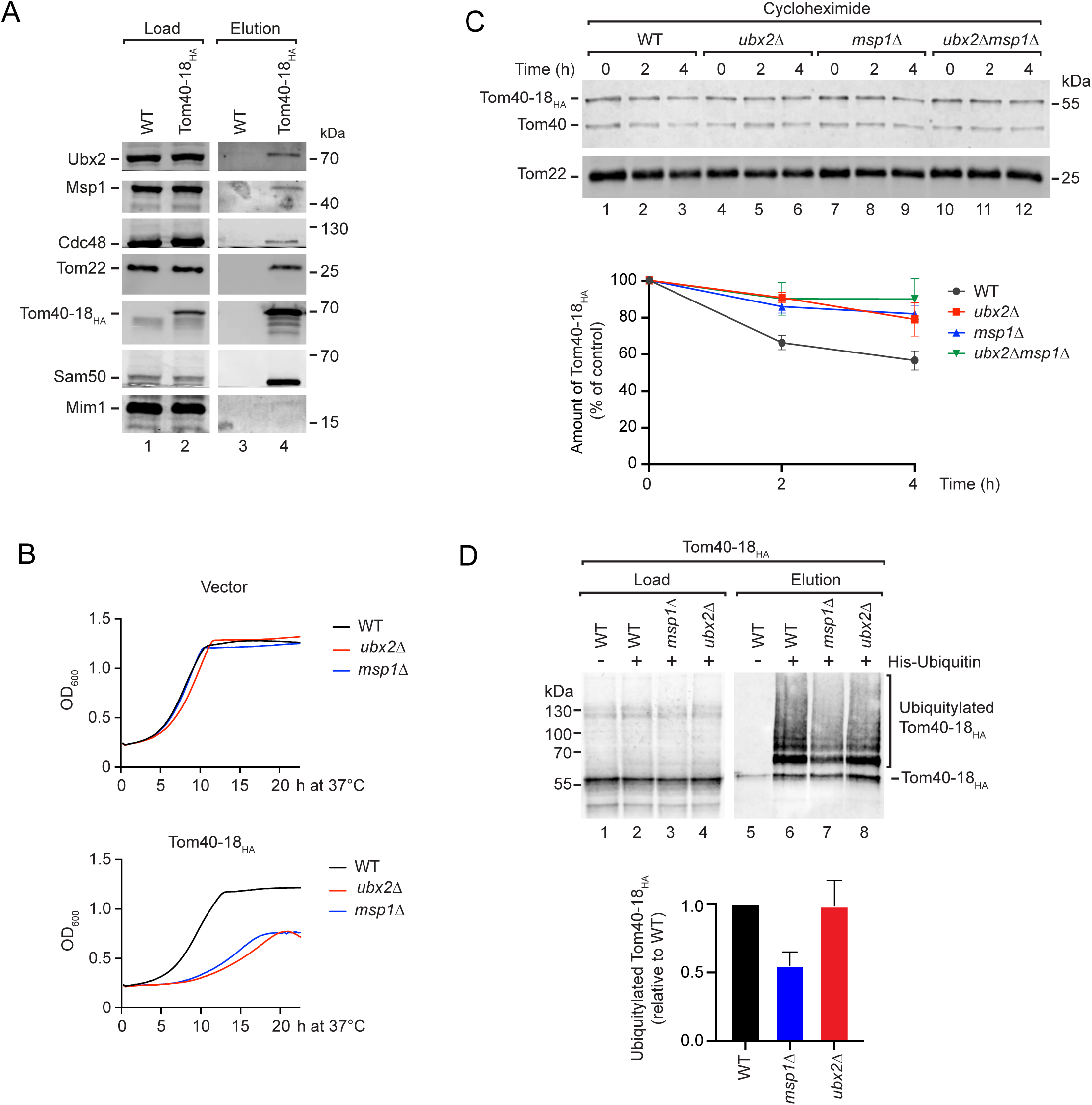
Msp1 and Ubx2 remove translocation-arrested Tom40-18_HA_. **(A)** Total cell extracts from wild-type (WT), Tom40_HA_ and Tom40-18_HA_ were subjected to affinity purification. Load (1%) and elution (100%) fractions were analyzed by SDS-PAGE and immunodetection with the indicated antisera. **(B)** Growth analysis of the indicated strains containing either the empty plasmid (upper panel) or a plasmid expressing *TOM40-18_HA_* (lower panel) in selective medium containing glucose as carbon source at 37°C. The optical density at 600 nm (OD_600_) was determined over time. **(C)** Upper panel, WT, *ubx2Δ*, *msp1Δ* and *ubx2Δ msp1Δ* cells expressing *TOM40-18HA* were treated with cycloheximide (CHX) for the indicated time periods. Cell extracts were analyzed by SDS-PAGE and immunodetection with the indicated antisera. Lower panel, quantifications of the Tom40-18_HA_ levels in the experiment shown in the upper panel. Depicted are mean values of four independent experiments with their S.E.M.. The time point 0 of each strain was set to 1 (control). **(D)** Upper panel: WT, *ubx2Δ* and *msp1Δ* cells expressing *TOM40-18HA* expressing His-tagged ubiquitin were subjected to affinity purification under denaturing conditions. Load (1%) and elution (100%) fractions were analyzed by SDS-PAGE and immunodetection with the indicated antisera. Lower Panel: Quantification of ubiquitylated Tom40-18_HA_ normalized to total ubiquitin signal. Depicted are mean values of three independent experiments with their S.E.M.. WT was set to 100% (control).

### Tom40-18HA is sorted to MitoStores

Unassembled Tom40-18_HA_ is extracted from the outer membrane through the coordinated action of the AAA ATPases Msp1 and Ubx2/Cdc48 to promote its proteasomal degradation. Non-imported mitochondrial precursor proteins can accumulate in distinct cellular compartments and protein deposits like the MitoStores ^53,55,56^. We therefore determined the cellular localization of Tom40-18_HA_ and Tom40_HA_ by immunofluorescence (Figures 4A, B). We found that Tom40-18_HA_ is sorted more frequently to protein deposits than Tom40_HA_ (Figure 4A). When cells were shifted to 37°C to challenge the proteostasis network, we detected Tom40-18_HA_ foci in almost all cells (Figure 4B). This observation suggests that Tom40-18_HA_ is transferred into protein deposits. To obtain an independent line of evidence, we lysed the cells with the mild detergent digitonin and separated soluble and insoluble proteins by centrifugation. Consistent with sequestration in protein deposits, the majority of Tom40-18_HA_ was recovered in the pellet fraction, whereas wild-type Tom40 was mainly detected in the soluble fraction (Supplemental Figure 2A). We wondered whether Tom40-18_HA_ sequestered in protein deposits is degraded by the proteasome. Upon cycloheximide treatment, the aggregated fraction of Tom40-18_HA_ decreased over time, and this turnover was blocked by treatment of the cells with the proteasomal inhibitor MG132 (Supplemental Figure 2B). We concluded that extracted Tom40-18_HA_ is temporally stored in protein deposits before its degradation by the proteasome.

**Figure 4.**
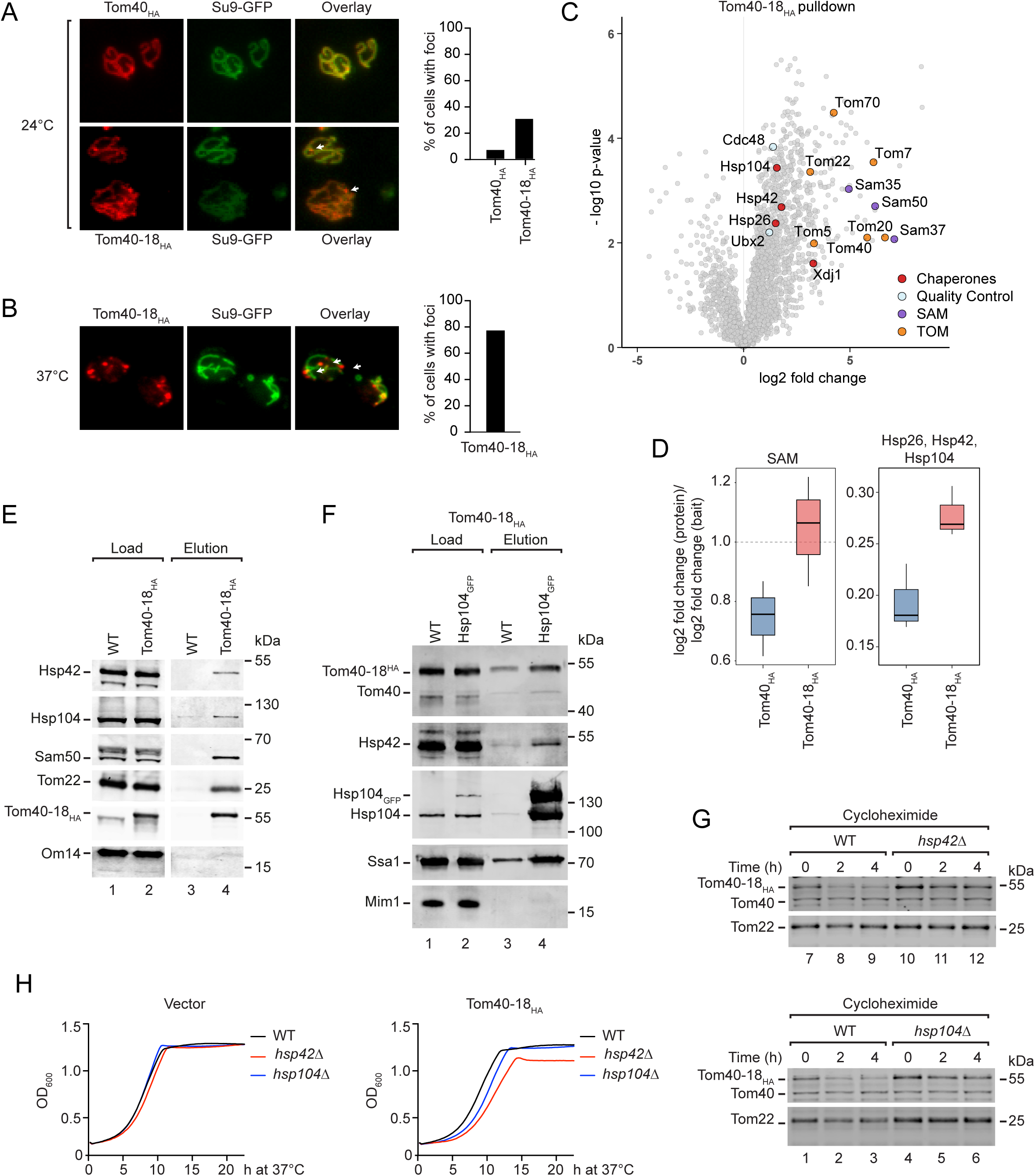
MitoStores facilitate the degradation of Tom40-18_HA_. **(A)** and **(B)** Right panel, Immunofluorescence of wild-type (WT) (A) or *pdr5*Δ (B) cells expressing TOM40-HA (A) and *TOM40-18HA* (A) and (B) and *Su9-GFP*, which were grown at 24°C (A) or shifted to 37°C (B). Tom40_HA_ and Tom40-18_HA_ were detected with specific antisera against the HA-tag. Arrows mark Tom40-18_HA_ present in protein deposits outside mitochondria. Left, quantification of cells containing Tom40_HA_ or Tom40-18_HA_-positive protein deposits. **(C)** WT cells with empty plasmid or expressing *TOM40-18_HA_* were lysed and subjected to affinity purification. The eluates were analyzed by mass spectrometry. Data are mean of four biological replicates. Statistical significance was determined via two-sided Welch’s t-tests with Benjamini–Hochberg correction. TOM, SAM, quality control factors and molecular chaperones are highlighted. **(D)** Ratio of co-purified chaperones and SAM subunits relative to the bait proteins Tom40_HA_ and Tom40-18_HA_ detected by mass spectrometry. Boxplots show the median, interquartile range, and whiskers. Corresponding group-level summary statistics, including mean and standard deviation are provided in Supplementary Table S3. **(E)** WT and Tom40-18_HA_ cells were lysed and subjected to affinity purification. Load (1%) and elution (100%) fractions were analyzed by SDS-PAGE and immunodetection with the indicated antisera. **(F)** WT and Hsp104_GFP_ cells expressing *TOM40-18HA* were lysed and subjected to affinity purification. Load (0.5%) and elution (100%) fractions were analyzed by SDS-PAGE and immunodetection with the indicated antisera. **(G)** WT, *hsp42Δ* and *hsp104Δ* cells expressing *TOM40-18HA* were treated with cycloheximide (CHX) for the indicated time periods. Cell extracts were analyzed by SDS-PAGE and immunodetection. **(H)** Growth of WT, *hsp42Δ* and *hsp104Δ* cells containing the empty vector (left panel) or expressing *TOM40-18HA* (right panel) at 37°C with glucose as carbon source. The optical density at 600nm (OD_600_) was determined at different time points.

Deposition of proteins into cytosolic inclusions such as MitoStores is mediated by small heat shock proteins like Hsp42 and Hsp26 ^55,56,73–76^, while their extraction depends on the hexameric AAA ATPase Hsp104 ^74–77^. Non-imported precursor proteins have been detected in such Hsp42-containing protein deposits, termed MitoStores ^56^. To investigate whether Tom40-18_HA_ is stored in MitoStores, we performed affinity purification of Tom40-18_HA_ and analyzed the elution fraction by mass spectrometry. Beside TOM and SAM components, Hsp104, Hsp42 and Hsp26 were detected in the elution fraction (Figure 4C, Supplemental Table 1). We next asked whether association of these chaperones with Tom40-18_HA_ was increased compared to their binding to Tom40_HA_. We compared the co-purification ratios of chaperones and SAM subunits recovered with Tom40-18HA versus Tom40_HA_ (Supplemental Fig. 2C and Supplemental Table 2). This analysis revealed increased association of both SAM subunits and the MitoStore-associated chaperones Hsp104, Hsp42 and Hsp26 with Tom40-18_HA_ (Figure 4D). We confirmed the association of Hsp104 and Hsp42 by Western blotting (Figure 4E, lane 4). Conversely, Tom40-18_HA_ can be co-purified along with GFP-tagged Hsp104 (Figure 4F, lane 4). We conclude that extracted Tom40-18_HA_ is deposited in MitoStores.

To understand why Tom40-18_HA_ accumulates in MitoStores, we asked whether sequestration in these deposits promotes its degradation. Formation of protein deposits organized by small heat shock proteins such as Hsp42 is generally enhanced when proteasomal capacity is challenged by excess client proteins ^56,74,75^. Hsp42 is a core component of the MitoStores and required for the formation of these protein deposits ^56,73–75^. Co-aggregation of substrates with Hsp42 allows efficient recovery, while aggregates formed in the absence of Hsp42 are more resistant to disaggregation ^76,78^. In our protein stability assay, the degradation of Tom40-18_HA_ is impaired in the absence of Hsp42 (Figure 4G), indicating that its ordered deposition in MitoStores supports its efficient degradation. The hexameric AAA ATPase Hsp104 disaggregates proteins from the MitoStores ^56^. Tom40-18_HA_ was stabilized in the absence of Hsp104 (Figure 4G), revealing that its Hsp104-mediated disaggregation delivers the Tom40-18_HA_ for proteasomal degradation. Formation of protein deposits such as MitoStores protects cellular proteostasis by sequestration of aberrant proteins ^56,78^. Consistent with this, cells lacking Hsp42 showed increased sensitivity to the expression of Tom40-18_HA_, whereas loss of Hsp104 did not cause a comparable growth defect under these conditions (Figure 4H). We conclude that after extraction from mitochondria Tom40-18_HA_ is stored in MitoStores to limit its proteotoxicity and to facilitates its ordered degradation by the proteasome.

## DISCUSSION

We report here the degradation pathway of the SAM-stalled β-barrel protein, Tom40-18_HA_. Extraction of this protein from the outer membrane is mediated by two AAA ATPases. The outer membrane-embedded AAA ATPase Msp1 promotes an initial extraction of Tom40-18_HA_ to facilitate its ubiquitylation. Ubx2 can bind ubiquitylated substrates via its UBA-domain and recruits Cdc48 via UBX domain ^46,71,72^. Cdc48 completes extraction of Tom40-18_HA_ from the mitochondrial outer membrane. Non-mitochondrial Tom40-18_HA_ can be deposited into MitoStores. Thereby, the unorganized accumulation of this hydrophobic Tom40-18_HA_ is diminished to minimize the proteotoxic stress in the cell. Hsp104 disaggregates Tom40-18_HA_ to facilitate its proteasomal degradation. We conclude that Tom40-18_HA_ is degraded via a multistep mechanism that involves one mitochondrial and two cytosolic AAA ATPases.

Sequestration of proteins into inclusions protects cells from an overload with damaged or incorrectly sorted proteins, which would sequester general protein quality control components and thereby compromise cellular proteostasis ^78,79^. Likewise, the ordered sequestration of damaged β-barrel proteins into MitoStores prior to disaggregation and proteasomal turnover might prevent these proteins from undergoing spurious interactions explaining the increased toxicity in cells lacking Hsp42. Newly translated β-barrel proteins associate with Hsp104 and Hsp42 in yeast translational extract ^15^, supporting our finding that MitoStores sequester hydrophobic β-barrel proteins when they are outside mitochondria. MitoStores were described as a temporary storage site for precursor proteins under proteotoxic stress conditions, which are eventually imported after the stress is relieved ^56^. Here, we uncovered that MitoStores can also store damaged β-barrel proteins to facilitate their degradation by the proteasome.

Protein quality control mechanisms are crucial for removing non-imported precursor proteins from mitochondria and the cell. So far, molecular mechanisms that extracts translocation-stalled precursor proteins have only been assigned for presequence-containing proteins ^46,58^. The molecular mechanisms that retro-translocate the other 30% of mitochondrial precursor proteins remain unknown. We report here that factors such as Msp1 and Ubx2 also retro-translocate β-barrel proteins, revealing their central role in mitochondrial proteostasis.

## SUPPLEMENTAL INFORMATION

The supplemental information includes two figures and two tables.

## ACKNOWLEDGEMENTS

Work in this study has also been performed in partial fulfilments of the requirements for the doctoral thesis of E.T.K. and L.C.S.. We thank Ralph Mahlberg and Hannah Scheuch for expert technical assistance. This study was supported by the Deutsche Forschungsgemeinschaft (DFG) (SFB 1218 B11 project ID 269925409, BE 4679/9-1 project ID 528247081, priority program SPP 2453 BE 4679/11-1 project ID 541555098, to T.B.; BR 6283/5-1 project ID 529716110, BR 6283/6-1 project ID 541596792, to F.d.B.) The mass spectrometer of the Core Facility “Analytical Proteomics”, University of Bonn, was funded by the DFG (project ID 386936527).

## AUTHOR CONTRIBUTIONS

Author contributions: E. T. Kämmerer, N. Okon, N. Limbach, E. Scifo, J. Hinnenkamp, L. Calvo Santos, M.H. Schuler, G. Schmidt, and F. Eiding performed the experiments and analyzed the data together with F. den Brave and T. Becker. T. Becker and F. den Brave designed and supervised the project; T. Becker, and F. den Brave designed and prepared the figures and wrote the manuscript; all authors discussed results from the experiments and commented on the manuscript.

## DECLARATION OF INTERESTS

The authors declare no competing interests.

## STAR METHODS

### LEAD CONTACT AND MATERIALS AVAILABILITY

Further information and requests for resources and reagents should be directed to and will be fulfilled by the Lead contact, Thomas Becker.

### EXPERIMENTAL MODEL AND SUBJECT DETAILS

Bakeŕs yeast *Saccharomyces cerevisiae* was used as model organism. All yeast strains have the genetic background of either BY4741 or YPH499. Information about the strains and their respective genotypes can be found in the KEY RESOURCES TABLE.

### METHOD DETAILS

#### Yeast strains and growth conditions

The wild-type yeast strains YPH499 and BY4741 have been described^80,81^. The *TOM40-18* mutant allele was generated by error prone PCR. It contains several point mutations (L55S, Y79H, F98S, A118T, N130S, N142S, T220I, S223C, R261S, I344T) that are distributed along the entire sequence. The genetic information of *TOM40-18* was inserted into a pFL39 plasmid and transferred into a Tom40 shuffle strain, in which the chromosomal open reading frame of *TOM40* was deleted using an *ADE2* selection marker. A wild-type copy of TOM40 was encoded by a pYEP352 plasmid that contains a URA3 marker. After transformation with pFL39-*TOM40-18* the plasmid encoding wild-type *TOM40* was removed by growth on 5-FOA-containing medium ^60^. To allow detection of both the mutant and the wild-type Tom40, the genetic information for a triple hemagglutinin-tag was inserted after the open reading frame of *TOM40-18* and before the STOP codon and subcloned into the p426 pADH vector. To study the degradation of the Tom40-18_HA_, p426 pADH-*TOM40-18-HA* was transformed into various yeast mutant strains (Key Resources Table). Cells were grown in selective medium (1.9 g/L selective medium lacking Uracil (Formedium), 6.7 g/L yeast-nitrogen-base with ammonium sulfate) using 2% (w/v) glucose as carbon source at 30°C or 37°C or were shifted from 24°to 37°C for 4 h when indicated. To determine the growth phenotype, the optical density at 600 nm of a cell culture was recorded using spectrostar nano plate reader (BMG labtech).

#### Isolation of mitochondria

Mitochondria were isolated by differential centrifugation following established protocols ^82^. Yeast cells were grown until the early logarithmic growth phase. Subsequently, cells were isolated (3,000 xg, 20°C, 5 min) and washed with water. Cells were treated with 0.1 M Tris pH 9.4 and 10 mM DTT for 30 min at 30°C under constant shaking. Cells were re-isolated and incubated with zymolyase (3 mg/ml) in zymolyase buffer for 45 min at 30°C. Cells were collected by centrifugation (3,000xg, 5 min, 20°C) and opened by homogenization in homogenization buffer. After removal of cell debris, mitochondria were isolated and washed with SEM buffer (250 mM sucrose, 10 mM MOPS, 1 mM EDTA). Mitochondria were resuspended in SEM buffer according to 10 mg/ml protein concentration, aliquoted and shock frozen in liquid nitrogen. Mitochondria were stored at -80°C until use.

#### *In vitro* protein import into isolated mitochondria

For protein import studies, we produced ^35^S-labelled proteins using a cell-free translation system based on reticulocyte lysate. We used as template the open reading frame of the genes of interest and placed it in a pGEM4z vector, which contains a strong SP6 promoter and the genetic information for a ribosome binding site. The plasmid encoding for the gene of interest was used for *in vitro* translation using the TNT kit (TNT Quick Coupled Transcription and Translation kit (Promega)) to incorporate [^35^S]methionine. In the import reaction, the radio-labelled precursor protein was incubated with isolated mitochondria in import buffer (3% (w/v) BSA, 250 mM sucrose, 80 mM KCl, 5 mM MgCl_2_, 2 mM KH_2_PO_4_, 5 mM methionine, 10 mM MOPS-KOH pH 7.2, 12 mM creatine phosphate, 0,1 mg/mL creatine kinase, 2 mM ATP, 2 mM NADH) for different time periods. The import of Atp2 and Su9-DHFR was blocked by depletion of the membrane potential with AVO mix (8 mM antimycin,1 mM valinomycin, 20 mM oligomycin), whereas the import of Tom40 and Por1 was stopped by transfer on ice. In the case of import of Atp2 and Su9-DHFR, mitochondria were treated with proteinase K (50µg/ml) in import buffer for 15 min on ice. Subsequently, the protease activity was inhibited by incubation with 0.2 M (final concentration) phenylmethylsulfonyl fluoride (PMSF) for 10 min on ice. Mitochondria were re-isolated and washed with SEM buffer. The mitochondrial pellets were lysed with 1% (w/v) SDS in Laemmli buffer. The import of Tom40 and Por1 was analyzed by blue native electrophoresis. Here, mitochondria were lysed with 1% (w/v) digitonin in lysis buffer (50 mM NaCl, 20 mM Tris/HCl pH 7.4, 0.1 mM EDTA, 10% (v/v) glycerol) for 15 min on ice. Under these conditions, mitochondrial protein complexes remain intact. After removal of non-solubilized material, the protein complexes were separated by blue native electrophoresis. To detect ^35^S-labeled proteins, we exposed the gels of import experiments to storage phosphor screens and detected the signal with a Typhoon FLA7000 and a Fuji FLA9000 phosphoimager.

#### Protein stability assay

Yeast cultures were grown in selective medium lacking uracil at 24°C until early logarithmic growth phase. Cells were diluted in fresh medium to an optical density (OD_600_) of 1.0 and shifted for another 3 h to 37°C. Protein stability was monitored over time after blocking protein synthesis by addition of 200 µg/mL cycloheximide to the growth medium. To confirm that cycloheximide treatment inhibits the growth of cells, the optical density (OD_600_) was measured before and after the experiment. For proteasomal inhibition *pdr5Δ* cells were treated with 100 µM MG132. After different time periods, cells were harvested by centrifugation (5 min, 20°C, 3,000 x g) and cell extracts were prepared followed published procedures ^83^.

#### Protein aggregation assays

Yeast cells corresponding to 15 OD_600_ were harvested (5 min, 20°C, 3,000 x g) and resuspended in fractionation buffer (300 mM NaCl, 100 mM HEPES-KOH pH 7.9, 1 % (v/v) Triton X-100, 1 x halt protease inhibitor, 20 mM NEM). The cell wall was opened by mechanic force using silica beads. After removal of cell debris and unbroken cells, aggregated proteins were collected by centrifugation (10 min, 4 °C, 17,000 x g). Total, supernatant and pellet fraction were analyzed by SDS-PAGE and immunodetection.

#### Protease accessibility assay

Isolated mitochondria were diluted in SEM buffer or EM buffer (10 mM MOPS, 1 mM EDTA). Subsequently, proteinase K was added to a final concentration of 20 µg/mL and incubated for 15 min on ice. Proteinase K activity was inhibited by the addition of 2 mM of PMSF for 10 min on ice. Mitochondria were re-isolated and washed with SEM buffer before analysis by SDS-PAGE and immunodetection.

#### Carbonate Extraction

Mitochondria were collected by centrifugation (10 min, 4 °C, 17,000 x g). The resulting pellet was resuspended in 0,1 M sodium carbonate, pH 11.5 and incubated on ice for 30 min. Subsequently, membrane and soluble fractions were separated by centrifugation (30 min, 4 °C, 125,000 x g). Proteins of the supernatant were precipitated with 14% (w/v) tri-chloroacetic acid. After 15 min of incubation on ice, the sample was centrifuged (15 min, 4 °C, 17,000 x g). The resulting pellet was washed in cold 100 % (v/v) acetone and pelleted (15 min, 4 °C, 17,000 x g). The samples were dried at 37 °C until the acetone was completely evaporated. The dried pellet was resuspended and analyzed by SDS-PAGE.

#### Affinity purification

We utilized the HA-tag fused to Tom40 or Tom40-18 for affinity purification from total cell extracts to determine their interaction partners. Cells were grown on selective medium until an early logarithmic growth phase, harvested and resuspended in lysis buffer (20 mM Tris-HCL pH 7.4, 50 mM NaCl, 2 mM Mg-Acetate, 10 % (w/v) glycerol, 1 mM PMSF, HALT protease inhibitor cocktail). Silica beads were added and the cell wall was disrupted by mechanic force at 4°C for 1 min. Cells were lysed with 0.5% (w/v) digitonin in lysis buffer and incubated for 30 min at 4°C under constant rotation. Insoluble material was removed by centrifugation (17,000 x g, 10 min, 4°C or 3000 x g, 10 min, 4°C for the detection of MitoStore-associated chaperones). The supernatant was incubated with anti-HA beads (Sigma-Aldrich) for 1.5 h at 4°C under constant rotation. Subsequently, beads were extensively washed with wash buffer (20 mM Tris-HCL pH 7.4, 50 mM NaCl, 2 mM Mg-Acetate, 10 % (w/v) glycerol, 1 mM PMSF, HALT protease inhibitor cocktail) containing 0.1% (w/v) digitonin and bound proteins were eluted by HU buffer (5 % SDS (w/v), 8 M Urea, 1 mM EDTA, 200 mM Tris-HCl pH 6.8, 0.025 (w/v) mM bromphenol blue). Pulldown of Hsp104-GFP was performed following the same protocol with the exception that cells were lysed with 1% (w/v) digitonin in the lysis buffer and 0.3% (w/v) digitonin was added to wash buffer. Pulldown was performed using magnetic GFP-trap beads (ChromoTek).

#### Sample preparation for mass spectrometry

Affinity purifications of Tom40-18_HA_ were performed in quadruplicate followed the procedure described above. Eluted proteins were denatured and loaded onto a SDS-PAGE. Gel lanes were cut into 3–4 slices for in-gel digestion. Gel pieces were destained, reduced (10 mM DTT, 20 min, 60°C), alkylated (50 mM iodoacetamide, 30 min, 20°C, dark), and digested overnight at 37°C with sequencing-grade trypsin. Peptides were extracted with acetonitrile, dried, reconstituted in 0.1% (v/v) formic acid, and 0.5–1 µg was injected per LC–MS/MS run.

#### NanoLC–MS/MS and data-dependent acquisition

Peptides were separated on a Dionex Ultimate 3000 RSLC nano-HPLC coupled to an Orbitrap Fusion Lumos (Thermo Fisher Scientific) in positive-ion mode using a self-packed C18 column (400 mm × 75 µm; 3 µm ReproSil-Pur 120 C18-AQ). Separation was performed at 300 nL/min using a gradient of 5–31% (v/v) solvent B over 75 min, followed by 31–50% (v/v) over 15 min (total runtime 122 min).

The instrument was operated in DDA mode with internal lock-mass calibration (m/z 445.12001). MS1 scans (330–1600 m/z) were acquired in the Orbitrap at 120,000 resolution (m/z 200). Precursors with charge states 2–7 were selected within a 2 s cycle using monoisotopic precursor selection and 60 s dynamic exclusion (10 ppm). MS2 spectra were generated by HCD (NCE 30) using a 2 m/z isolation window and Orbitrap detection at 7,500 or 15,000 resolution depending on precursor intensity.

#### Protein identification and quantification

Raw files were processed in Proteome Discoverer 3.2 using Mascot v2.8.1 against the *Saccharomyces cerevisiae* UniProt proteome (release 2025-06-18) supplemented with CRAPome contaminants. Searches used trypsin specificity (≤2 missed cleavages), 10 ppm precursor tolerance, and 0.02 Da fragment tolerance. Cysteine carbamidomethylation was set as fixed; methionine oxidation and protein N-terminal acetylation were variable. FDR was controlled at 1% at the PSM, peptide, and protein levels. LFQ was performed in Proteome Discoverer, retaining proteins identified by ≥2 unique peptides.

#### Quality control and statistical analysis

The performance of the mass spectrometer was monitored using BSA digest standards. Data quality was assessed by mass accuracy, chromatographic stability, and replicate reproducibility (Pearson r ≥ 0.95; CV ≤ 20%). Protein abundances were analyzed in R (version 4.5.1) after log2 transformation and Perseus-like missing-value imputation. Tom40 pulldown samples were normalized to the median Tom40 signal within each group. Proteins with ≥3 valid values in one group were retained. Differential enrichment was assessed using log2 fold change and two-sided Welch’s t-tests with Benjamini–Hochberg correction; significance thresholds were adjusted p < 0.05 and |LFC| ≥ 1.

#### Ubiquitylation assay

Ubiquitylation was analyzed following established protocol ^84^. We co-expressed plasmid-borne His-tagged ubiquitin ^59^ with Tom40-18_HA_ in wild-type and *msp1*Δ and *ubx2*Δ strains. The strains were grown in selective medium that lack both uracil and leucine and contains 2% (w/v) glucose as carbon source. Cells were harvested and resuspended in 6 M guanidine HCl, 100 mM NaH_2_PO_4_ and 10 mM Tris and mechanically ruptured using silica beads. 0.05% (v/v) Tween 20 and imidazole (20 mM final concentration) were added and samples were incubated for 20 min on ice. Insoluble material was removed by centrifugation (17,000 x g, 10 min, 4°C) and the supernatant was incubated with Ni-NTA agarose (Qiagen) for 1h 30 min at 4°C under constant rotation. Subsequently, the beads were washed with an excess volume of 8 M urea, 100 mM NaH_2_PO_4_ and 10 mM Tris. Proteins were eluted with HU buffer and analyzed by SDS-PAGE and immunoblotting.

#### Immunoblotting

For immunodetection, proteins were transferred onto polyvinylidene fluoride membrane (PVDF) (EMD Milipore) by semidry Western blotting. After washing with TBS buffer (20 mM Tris/HCl pH 7.4, 12.5 mM NaCl), the membranes were blocked with 5% (w/v) skimmed milk powder in TBS buffer or with RotiBlock (Roth) solution to minimize unspecific background signals. Subsequently, the membranes were washed with an excess amount of TBS buffer. We used different antisera to detect mitochondrial and proteostasis proteins by immunoblotting (KEY RESOURCES TABLE). We used two approaches to detect the proteins. First, enhanced chemilumnescence (ECL)-staining ^85^ was used to detect the immunosignals by Amersham Imager 680 (Cytiva). Second, we used secondary antibodies against rabbit or mouse coupled to fluorescent labels (IRDye 800CW or 680RD) and detected immunosignals using the Odyssey CLx Infrared Imaging System (Li-Cor). The obtained data were analyzed using Image Studio software (Li-Cor) and ImageJ software. The specificity of the immunosignals were controlled using cellular and mitochondrial lysates of deletion strains or strain containing tagged proteins. In deletion strains, the immunosignal was absent and in cells a expressing a tagged variant of the corresponding protein the immunosignal was shifted in size. We used Novex Sharp Pre-stained (Invitrogen) as molecular weight marker to determine the size of proteins on SDS-PAGE and the HMW Calibration kit for native electrophoresis as size marker for blue native electrophoresis.

### QUANTIFICATION AND STATISTICAL ANALYSIS

We used the freely available ImageJ software or ImageStudio (Li-COR) to process the obtained image files from Western blotting and autoradiography. We used Photoshop (Adobe) to process images and Illustrator (Adobe) to assemble the figures. When non-relevant bands were removed digitally, we separated the gel images. Statistical analysis of quantifications was performed using GraphPad Prism (GraphPad Software Inc., San Diego, USA).

### DATA AND CODE AVAILABILITY

The mass spectrometry data have been deposited to the ProteomeXchange Consortium via the jPOST ^86^ partner repository and are publicly available under the dataset identifiers PXD077794 and JPST004589. The data are available (http://proteomecentral.proteomexchange.org/) using the access key 1806. All other data supporting the reported findings are in this article or in the Supplemental Information.

### KEY RESOURCES TABLE

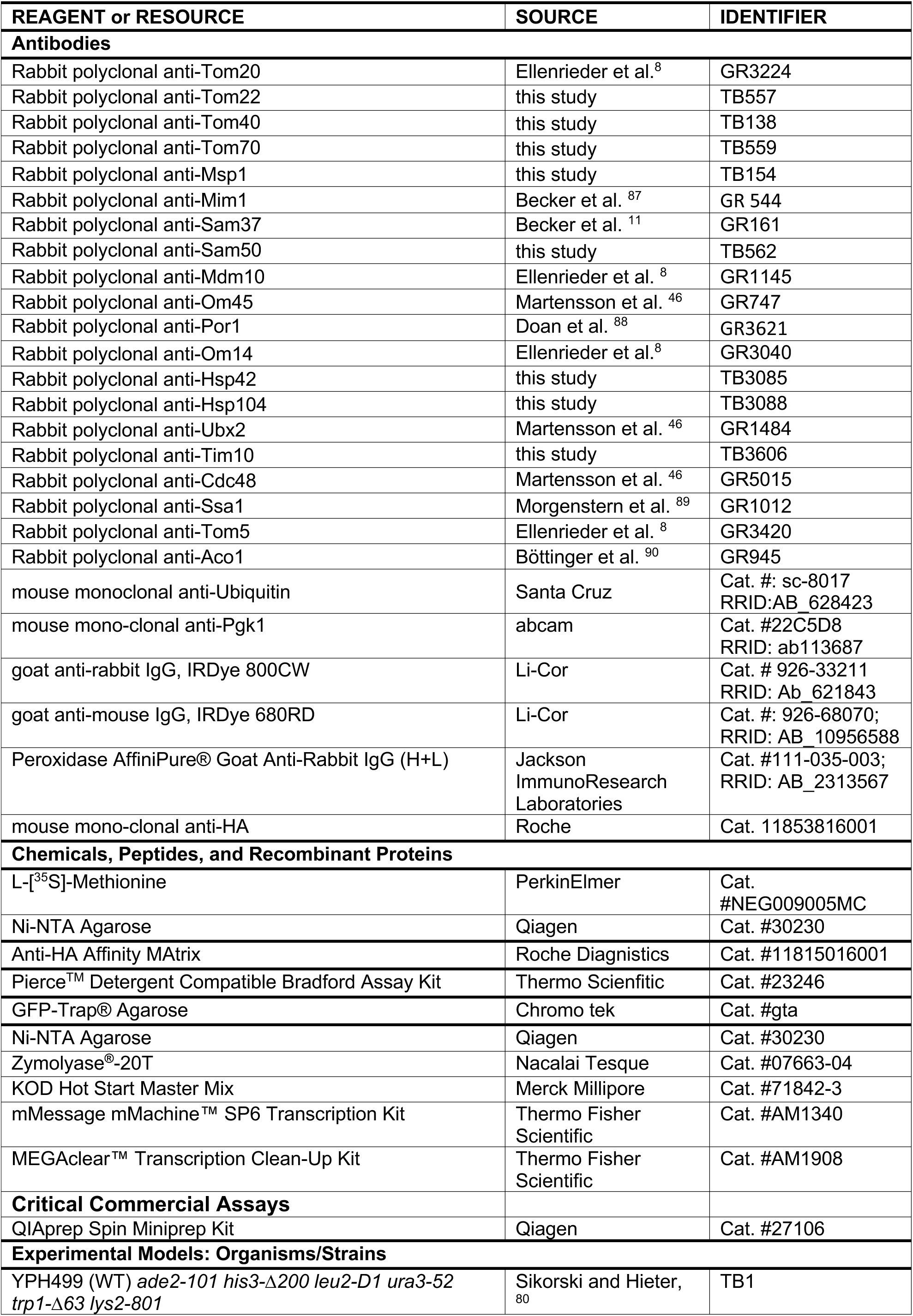
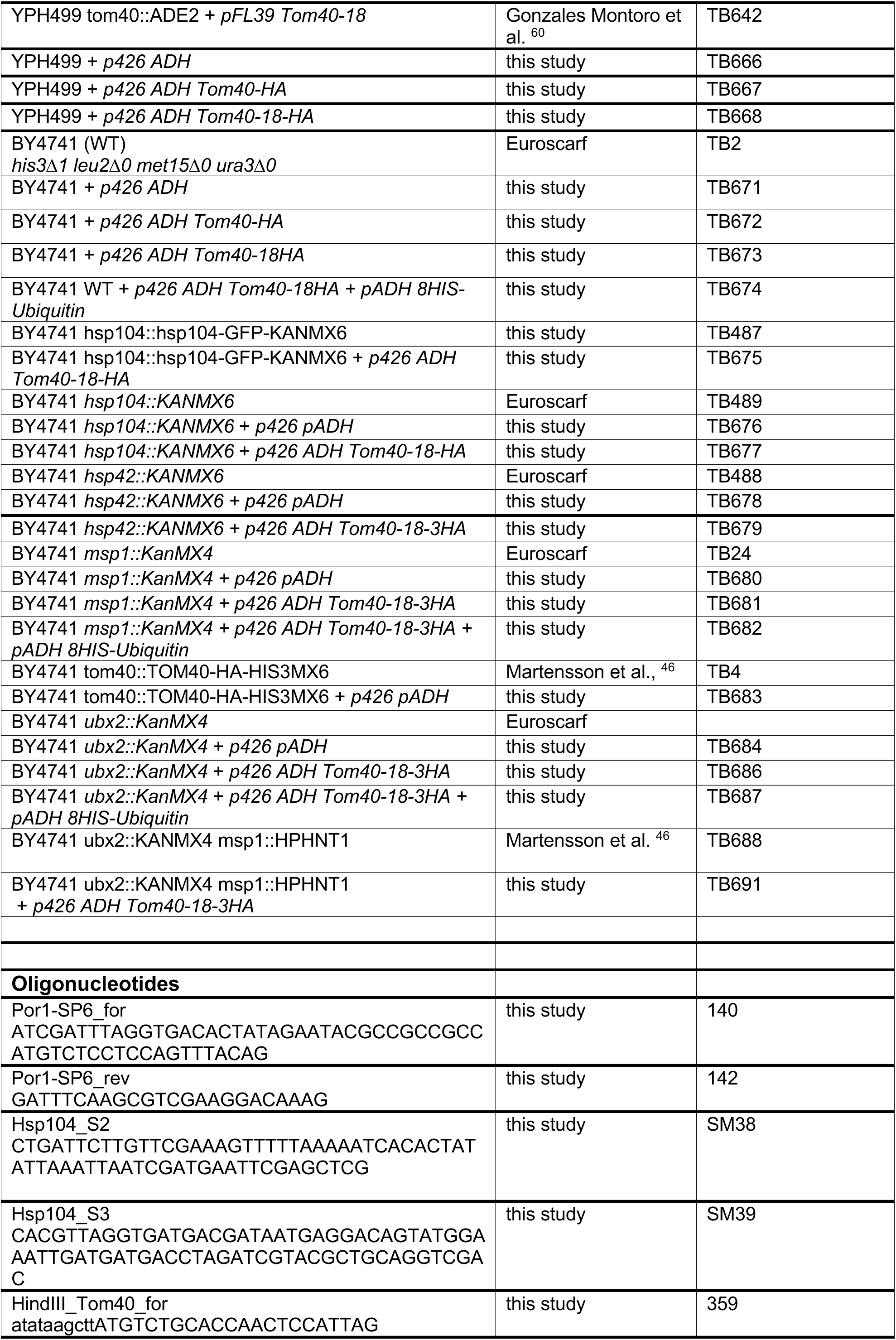
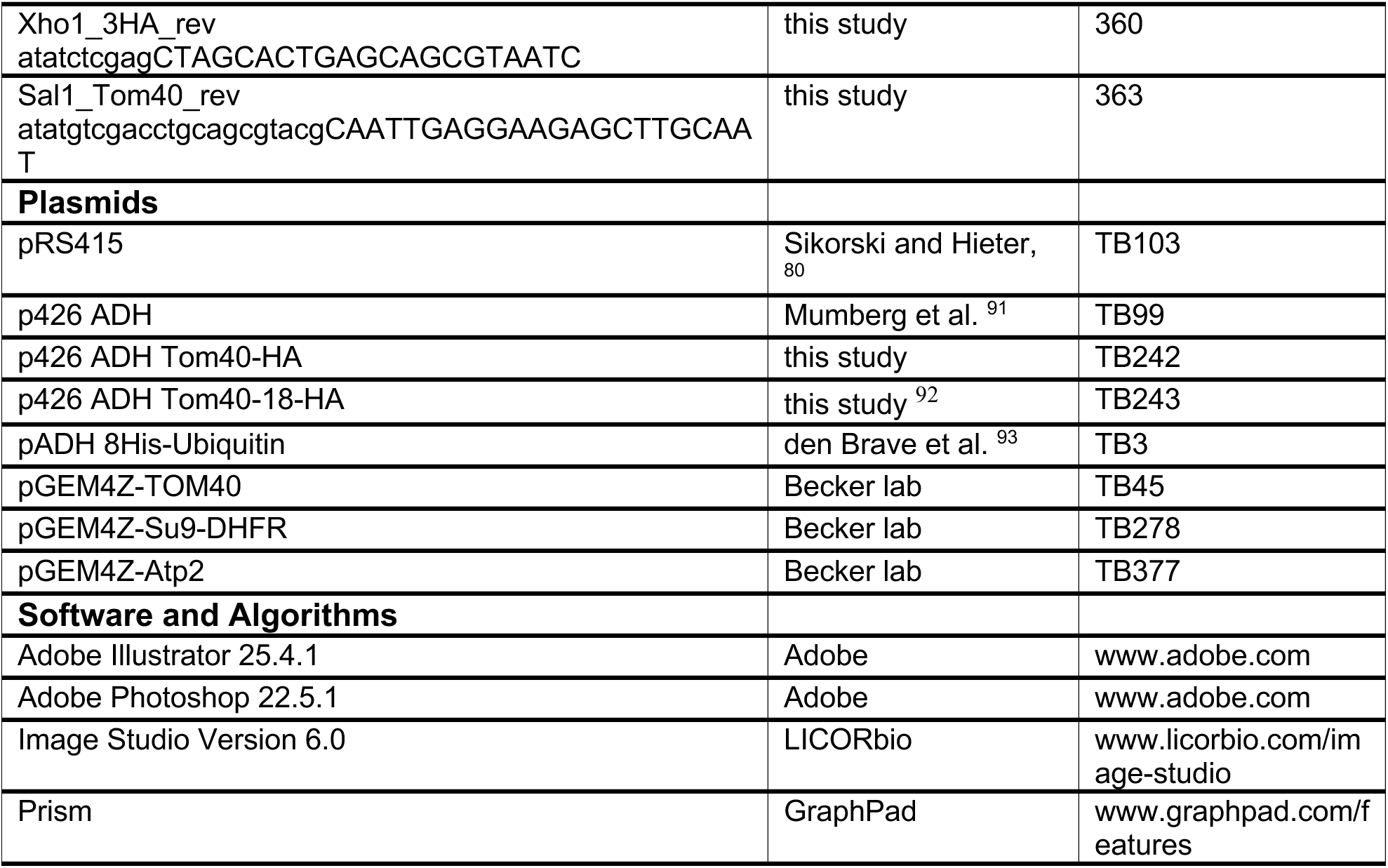

## Notes

### Competing Interest Statement

The authors have declared no competing interest.

