## Supplemental figures for "The multistep quality control of mitochondrial β-barrel protein import"

### FIGURE LEGENDS

**Figure S1. Characterization of Tom40-18 mutant strains. Related to Figure 1.**
**(A)** Proteins of wild-type (WT) and *tom40-18* mitochondria were analyzed by SDS-PAGE and immunodetection with the indicated antisera. **(B)** Serial dilutions of wild-type (WT) containing an empty vector or expressing $TOM40\text{-}18_{HA}$ were grown on selective medium containing glucose as carbon source at 37°C. **(C)** Immunofluorescence of WT cells expressing $TOM40_{HA}$ or *TOM40-18HA* and *Su9-GFP*, that were grown under permissive conditions. $Tom40\text{-}18_{HA}$ was detected with specific antisera against the HA-tag. **(D)** WT and *tom40-18HA* mitochondria were subjected to carbonate extraction. Pellet and supernatants were analyzed by SDS-PAGE and immunodetection with the indicated antisera. **(E)** WT and *tom40-18HA* mitochondria were treated under iso-osmotic or hypo-osmotic conditions with or without proteinase K. The samples were analyzed by SDS-PAGE and immunodetection with the indicated antisera.

**Figure S2. Tom40-18 is sequestered in protein deposits. Related to Figure 4.**
**(A)** Wild-type (WT) and $Tom40\text{-}18_{HA}$ cells (total, T) were lysed and separated into soluble (S) and pellet (P) fractions by centrifugation. Proteins were analyzed by SDS-PAGE and immunodetection with the indicated antisera. **(B)** *pdr5Δ* cells expressing $Tom40\text{-}18_{HA}$ cells were treated with the proteasomal inhibitor MG132 (total, T), were lysed and separated into soluble (S) and pellet (P) fractions by centrifugation. Proteins were analyzed by SDS-PAGE and immunodetection with the indicated antisera. **(C)** Wild-type cells with empty plasmid or expressing $TOM40_{HA}$ were lysed and subjected to affinity purification. The eluates were analyzed by mass spectrometry. Data are mean of four biological replicates. Statistical significance was determined via two-sided Welch's t-tests with Benjamini–Hochberg correction. TOM, SAM, quality control factors and molecular chaperones are highlighted.

### SUPPLEMENTAL TABLES

**Table S1.** List of proteins identified in the affinity purification via Tom40-18$_{HA}$ by quantitative MS.

**Table S2.** List of proteins identified in the affinity purification via Tom40$_{HA}$ by quantitative MS.

**Table S3.** Combined protein-level to bait values and statistics used for boxplot generation.

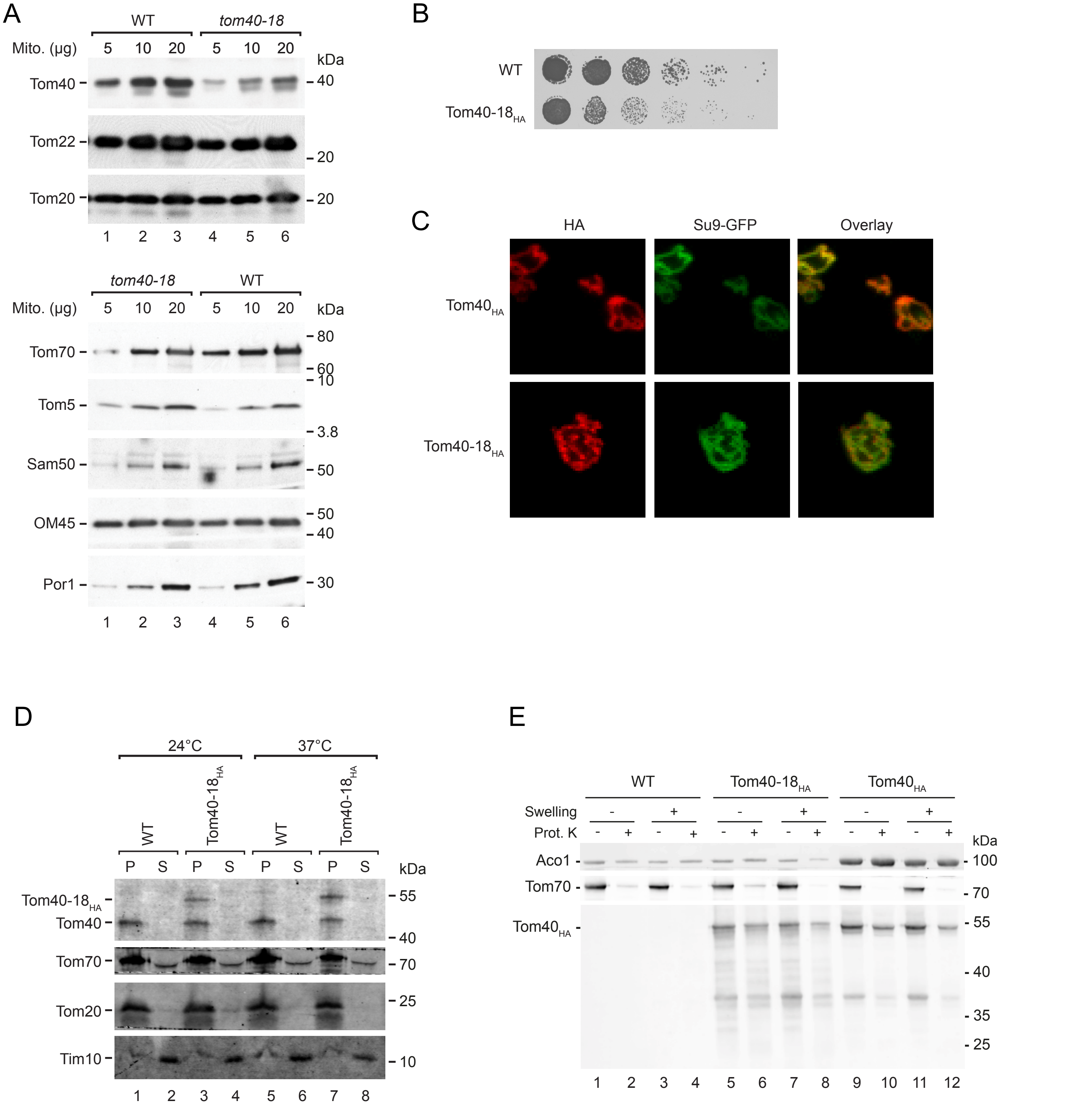


Supplemental Figure 1 - Kämmerer et al.

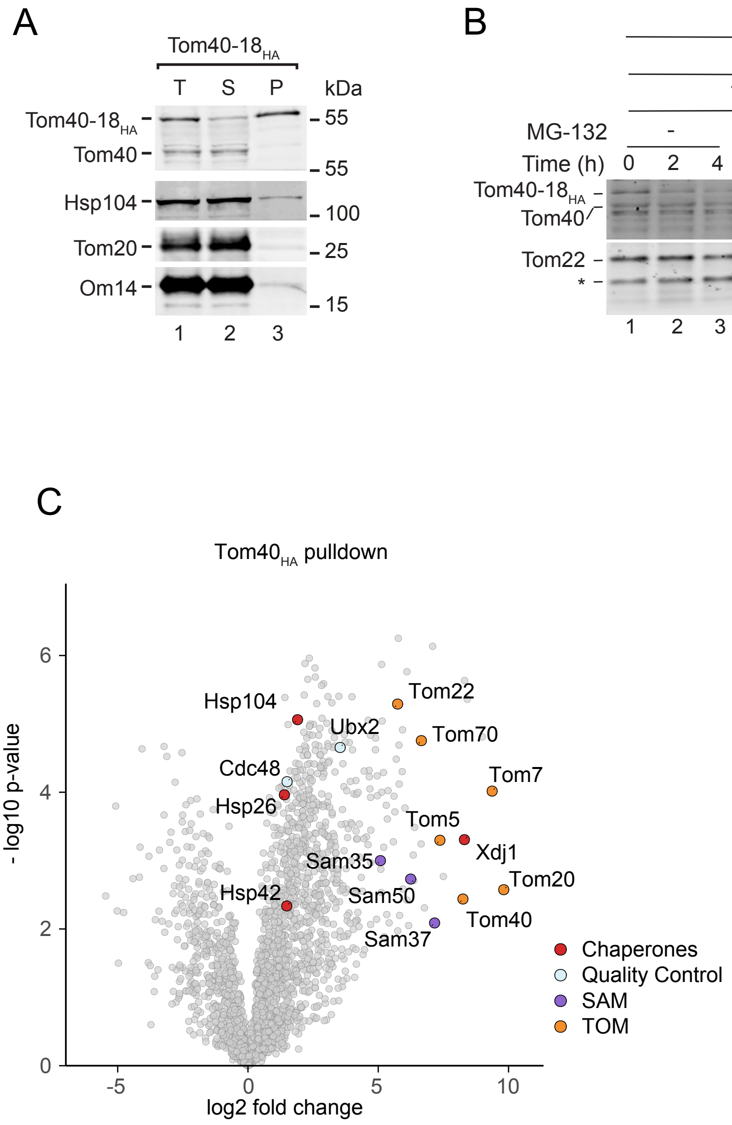


Supplemental Figure 2 - Kämmerer et al.
